# Overexpression of *magA* in *Acidithiobacillus ferrooxidans* increases magnetosome production and pyrite bioleaching

**DOI:** 10.64898/2026.05.26.727969

**Authors:** Heejung Jung, Sameera Abeyrathna, Zihang Su, Scott Banta

## Abstract

Acidithiobacillus ferrooxidans, a chemolithoautotrophic iron- and sulfur-oxidizing acidophile, is a key contributor to industrial-scale copper metal bioleaching. These cells naturally produce magnetosomes, and they may serve as an emerging platform for magnetosome bioproduction, as magnetotactic bacteria (MTB) are difficult to cultivate and to genetically modify. Here we manipulated the expression of the endogenous homologs to the magA and mamB genes in A. ferrooxidans, which are implicated in iron transport required for magnetosome synthesis. Modulation of mamB had no impact on cell behavior. Overexpression of magA increased magnetosome formation and magnetic responsiveness and theses effects were attenuated by CRISPRi knockdown of magA. The augmented magnetosome formation in the magA overexpression cells also led to enhanced bioleaching of pyrite, which is weakly paramagnetic, and this could be further enhanced by addition of an external magnetic field. These results confirm that magA plays a critical role in magnetosome formation in A. ferrooxidans and that magnetosome expression can be enhanced through genetic engineering. In addition, these results demonstrate the potential to improve metal sulfide bioleaching through the manipulation of genes involved in magnetosome formation.

**TOC Graphic:** 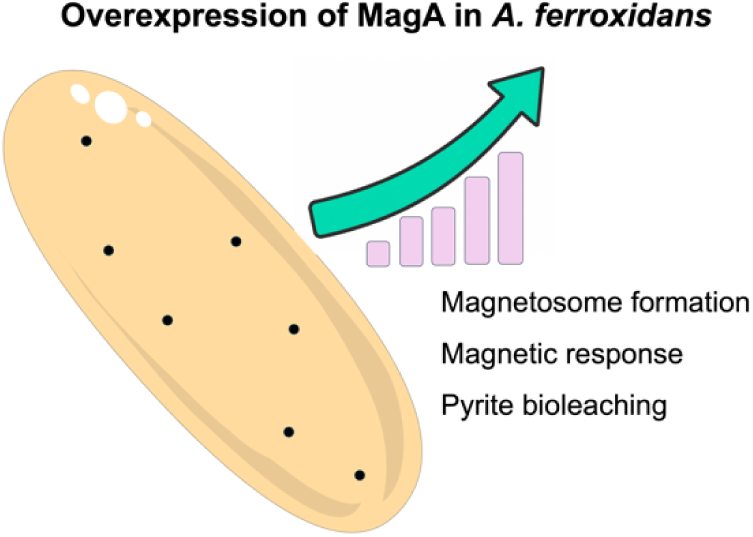

## Introduction

Magnetotactic bacteria (MTB) have the ability to synthesize magnetosomes, which are unique membrane-enclosed organelles that contain magnetic iron minerals, predominantly nanosized magnetite and greigite^1^. The exceptional properties of these bionanomaterials, including high purity, narrow size distribution, consistent crystal morphology, permanent ferrimagnetism, and excellent biocompatibility^2, 3^, have sparked substantial interest for a number of biotechnological applications. However, the difficult growth conditions required by MTB pose challenges for their mass cultivation on an industrial scale, and this hinders the elucidation of the molecular pathways involved in magnetosome biosynthesis. More recently, it has been discovered that magnetosomes can be synthesized by other metal oxidizers, including *Acidithiobacillus ferrooxidans.* This chemolithoautotrophic acidophile (thrives at pH <3)^4^ can perform dissimilatory iron oxidation or reduction and has some similar traits and growth conditions as MTB^3, 5^. Although the magnetosome yield in *A. ferrooxidans* is low, the relatively easier, aerobic cultivation conditions ^6^ present an alternative avenue for scalable magnetosome production^7^.

While investigations into magnetosome synthesis in *A. ferrooxidans* are still in the early stages, previous studies have characterized the physicochemical properties of magnetosomes^6, 8^ and identified growth conditions to enhance yields^9-12^. The mechanisms governing magnetosome synthesis in *A. ferrooxidans* remain largely unexplored, and they appear to differ from other MTB^13^. Recent studies have proposed models of magnetosome synthesis in the *A. ferrooxidan*s BYM strain via whole-genome sequencing^13^ and phenomics and transcriptomic analyses^14^. Magnetosome are predicted to be synthesized via membrane formation, iron uptake and transport, initiation of magnetite formation, followed by maturation of magnetosome particles. Although the involvement of several proteins in each step of magnetosome synthesis has been proposed, the specific functions of these proteins remain unconfirmed.

A previous bioinformatic analysis of the *A. ferrooxidans* ATCC 23270 genome identified ORF 1124 and ORF 2572, corresponding to AFE_1968 and AFE_0465, as candidate homologs of magnetotactic bacterial *magA* and *mamB*, respectively. These ORFs were reported to share 32% and 29% identity with *magA* and *mamB*, respectively, and were annotated as a potassium-efflux-system protein and a cation-efflux-family protein respectively. Liu et al. proposed that these proteins may participate in iron transport during magnetosome formation^10^. The MagA-like homolog, was annotated as a KefB/TrkA-domain-containing potassium-efflux-system protein and proposed to function as a proton-driven H^+^/Fe(II) antiporter. This proposed role was supported by Fe^2+^-responsive expression of MagA, increased TEM-visible intracellular particles under higher Fe^2+^ conditions, and enhanced magnetophoresis, although direct Fe(II) transport activity was not biochemically demonstrated^10^. In magnetotactic bacteria, MamB and MagA have been proposed to contribute to magnetosome-associated iron handling, although MamB has a more established role in magnetosome formation, while MagA function appears to be species-dependent^10, 15, 16^. Considering these two genes may support enhanced iron transport into magnetosomes, we hypothesized that modulating the expression of these genes could influence magnetosome formation in *A. ferrooxidans*. In this study, we genetically manipulated *A. ferrooxidans* to constitutively overexpress the native *magA* and *mamB* genes to attempt to enhance magnetosome formation. In addition, CRISPR interference (CRISPRi) ^17^ was used to knockdown *magA* and *mamB* expression to further explore the functional roles of these proteins in magnetosome biosynthesis. Our results demonstrate that manipulation of the *magA* gene can modulate magnetosome biosynthesis and that overexpression of *magA* not only increased magnetosome formation, but also led to enhanced bioleaching of pyrite.

## Materials and methods

### Strains, media, and chemicals

Strains used in this study include *E. coli* DH5a and *E. coli* DH10b purchased from NEB (Ipswich, MA), *E. coli* S17-1 ATCC 47055 and *A. ferrooxidans* ATCC 23270 obtained from ATCC (Manassas, Virginia). The modified AFM3 growth medium [2.4 g/L (NH_4_)_2_SO_4_, 0.1 g/L K_2_HPO_4_, 2.0 g/L MgSO_4_·7H_2_O, 5 mL/L Trace mineral solution (MD-TMS, ATCC), 1.92 g/L citric acid, 34.8 g/L FeSO_4_·7H_2_O, final pH of 1.8)] was used, in which the iron and nitrogen concentrations were previously optimized for magnetosome formation in *A. ferrooxidans* using a response surface methodology approach *^18^*. AFM1 medium (pH 1.8) contains 0.8 g/L (NH_4_)_2_SO_4_, 0.1 g/L K_2_HPO_4_, 2.0 g/L MgSO_4_·7H_2_O, 5 ml/L MD-TMS; and 20 g/L FeSO_4_·7H_2_O,. The SM4 medium (pH 5.0) consists of 0.8 g/L (NH_4_)_2_SO_4_, 0.1 g/L K_2_HPO_4_, 2.0 g/L MgSO_4_·7H_2_O 5 ml/L MD-TMS, 0.19 g/L citric acid, 17.9 mg/L Fe_2_(SO_4_)_3_, 40 mg/L leucine, 19 mg/L diaminopimelic acid, and dispersed 1 g/L sulfur, ^19^ (#S789400; Toronto Research Chemicals, Toronto, ON). The pH of the media was adjusted by adding sulfuric acid and the media was filtered through a 0.2 µm pore size PVDF filter (Thermo Fisher Scientific, Waltham, MA) before use. Dispersed sulfur^19^ was added to SM4 medium after filtration.

Enzymes and reagents for DNA manipulation were obtained from NEB, and oligonucleotides were purchased from Integrated DNA Technologies (Coralville, Iowa). All strains, plasmids, and oligonucleotides used in this study are presented in Tables S1 and S2.

Rare earth (neodymium) permanent square magnets (1.16 x 1.16 x 0.08 in) were from DIYMAG. Pyrite (FeS_2_, Cat# 77817) was purchased from Sigma-Aldrich (St. Louis, MO), and chalcopyrite (CuFeS_2_) concentrate (24.8% Cu, 27.2% Fe, and 30.7% S) was provided by Freeport-McMoRan (Phoenix, AZ), as previously described^20^. All chemicals were purchased from Sigma-Aldrich (St. Louis, MO), unless otherwise noted.

### Cell culture and harvest

All *A. ferrooxidans* strains were grown in 100 mL of modified AFM3 growth with an initial optical density measured at 600 nm (OD_600_) of 0.001, which corresponds to a cell density of 8.3 × 10^6^ cells/mL^21^. The cells were incubated in a shaking incubator (30 and 140 rpm). The cells were harvested by centrifugation at 5,000 *g* for 7 min. The harvested cells were repeatedly washed by resuspending with the basal salt solution (0.8 g/L (NH_4_)_2_SO_4_, 0.1 g/L K_2_HPO_4_, 2.0 g/L MgSO_4_·7H_2_O, 5 mL/L MD-TMS; pH 1.8) followed by centrifugation at 17,000 *g* for 1 min, unless otherwise noted.

### Plasmid construction and genetic modification

Genomic DNA of *A. ferrooxidans* was extracted using the NucleoSpin Tissue kit (Takara Bio, Mountain View, CA). To make pMagA and pMamB (Table S1), the sequences of putative *magA* and *mamB* genes, encoding putative MagA (AFE_1968) and MamB (AFE_0465) proteins in *A. ferrooxidans*, were amplified with designed primer sets (Table S2), then purified from the gel. The empty pJRD215 vector, pYI39 ^22^ was digested with KpnI restriction enzyme and treated with shrimp alkaline phosphatase (NEB) for dephosphorylation of the digested plasmid DNA. Each PCR amplicon was then combined with the digested pYI39 by NEBuilder HiFi DNA Assembly, as per the instructions of the manufacturer. The constructs were transformed into *E. coli* DH5α for sequence verification. To make pdMagA and pdMamB (Table S1), two guide RNA sequences for each target gene (Table S3) were inserted into dCas12a/CRISPR system constructed on pJRD215 vector (pJRD_dCas12a) ^17^using Q5 site-directed mutagenesis (NEB) with designed primer sets (Table S2). The plasmid constructs were transformed into *E. coli* DH10b to verify sequence. After sequence verification, all four plasmids were transformed into *E. coli* S17-1, then conjugally transferred to *A. ferrooxidans* by filter mating, as previously described^23^. After the conjugation, the obtained transconjugant colonies were initially grown in AFM1 medium, then isolated in low-iron, higher-pH, sulfur-based SM4 medium supplemented with 250 μg/ml kanamycin over three passages. After confirmation of overexpression or downregulation of the target genes at transcriptional level, the isolated strains were referred to as MagA, MamB, dMagA, and dMamB cells, respectively.

The nucleotide sequences of pMagA, pMamB, pdMagA, and pdMamB are deposited in Genbank with accession numbers of PP597399–597402.

### Transcriptional gene expression

Total RNA extraction of both wild type and engineered cells was performed by RNeasy Mini kit (Qiagen, USA), and reverse transcription to cDNA was carried out using QuantiTech Reverse Transcription Kit (Qiagen), according to the protocols. The transcriptional expressions of *magA* and *mamB* genes encoding MagA and MamB proteins, respectively, were measured by QIAcuity Digital PCR (dPCR) System (Qiagen, USA). The primer information used for dPCR analysis is present in Table S2.

### Quantification of intracellular magnetite

The functional effects of the overexpression or knockdown of *magA* and *mamB* on magnetosome formation in *A. ferrooxidans* were first examined by quantification of intracellular magnetite and total iron content. Both wild type and engineered cells were initially grown in modified AFM3 medium and harvested as described above. The final cell resuspension was split to measure intracellular magnetite and total iron, and centrifuged (17,000 g for 1 min). One portion of the cell pellet was resuspended in 10% HNO and digested at 80 °C for 30 min. The digested sample was centrifuged at 17,000 × g for 1 min to remove insoluble debris, and the supernatant was diluted with ultrapure water to adjust HNO to 1 % and subjected to an atomic absorption spectrometry (AAS; iCE 3300, Thermo Fisher Scientific, Waltham, MA), to measure the intracellular iron. Another portion of cell pellet was used for magnetite quantification via oxalate extraction method ^24, 25^ with slight modification. Briefly, cell pellets were resuspended in a FastBreak Cell Lysis Reagent (Promega, Madison) and incubated for 30 min at room temperature under shaking. After cell lysis, the supernatant was reacted with 0.1M oxalate solution (13.4 g/L sodium oxalate and 9 g/L oxalic acid, pH 2.5) (1:1, v/v) for 48 h in a shaking incubator (30 and 140 rpm). All solutions were filtered was filtered through 0.2 µm pore size PVDF filters (Thermo Fisher Scientific, Waltham, MA). Then, iron concentration extracted from magnetite was measured using AAS. Both magnetite and total iron concentrations were normalized by the cell density.

### Transmission electron microscopy (TEM)

To visualize the intracellular magnetosomes, cells were observed under TEM. The cells were grown in modified AFM3 medium in the absence or presence of external magnetic fields and incubated for 60 h prior to harvest. For the conditions with magnetic fields, a permanent neodymium magnet was attached to the bottom exterior of 250 mL Erlenmeyer flasks throughout the incubation. The cells were harvested by centrifugation at 5,000 *g* for 7 min. The harvested cells were then washed 3-5 times with the basal salt solution (0.8 g/L (NH_4_)_2_SO_4_, 0.1 g/L K_2_HPO_4_, 2.0 g/L MgSO_4_·7H_2_O, 5 mL/L MD-TMS; pH 1.8) at 17,000 *g* for 1 min. Cell resuspensions were applied to 200-mesh carbon film copper grids (Sigma-Aldrich) that had been plasma cleaned using a Harrick Plasma Cleaner (model PDC-001-HP) for 30 seconds at a low RF setting. Three microliters of sample were deposited onto each plasma-cleaned grid and incubated for 30 seconds. The excess sample was blotted using filter paper. Subsequently, 3 µL of ultrapure water was pipetted onto the grid and immediately blotted off. This washing step was repeated with 3 µL of ultrapure water one more time. After washing step, the grids were air-dried for approximately 3-6 h and imaged without chemical fixation by a FEI Talos F200X TEM.

### Modified whole-cell magnetic-response assay

A modified whole-cell magnetic-response assay was developed to determine whether the oxalate-soluble iron fraction was associated with magnetically responsive cell-associated material. The assay was adapted from the principle of the *C_mag_* light-scattering assay described by Schüler et al., in which magnetosome-containing cells align in an external magnetic field and produce a measurable change in optical scattering.^26^ In the present study, a simplified magnet-induced OD_600_ depletion assay was used.

A modified cuvette was prepared by placing two strong permanent neodymium magnets externally against opposite side walls of a BRAND^®^ 1.5 mL semi-micro cuvette. The magnets were positioned outside the sample chamber, granting a lateral magnetic field across the cuvette, perpendicular to the optical path of the spectrophotometer (Fig. S1).

The wild type and engineered cells with overexpression and knockdown of the *magA* gene were initially grown in modified AFM3 medium and harvested as described earlier. Wild type, MagA, MamB, and dMagA cell suspensions were normalized to an initial OD_600_ of 0.4. From each sample, 1.0 mL of normalized cell suspension was transferred either to the modified magnet-containing cuvette or to an identical control cuvette without magnets. Both cuvettes were incubated undisturbed for 10 min at room temperature. After incubation, OD_600_ was measured directly without disturbing the sample. The magnet-induced OD_600_ change was calculated as:

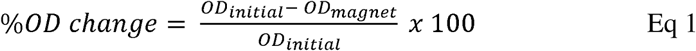

where OD_initial_ is the OD_600_ before incubation and OD_magnet_ is the OD_600_ after 10 min exposure in the magnet-containing cuvette. To correct for passive settling or nonspecific OD_600_ changes during the incubation period, a matched no-magnet control was included for each sample. Magnet-specific OD_600_ change was calculated as:

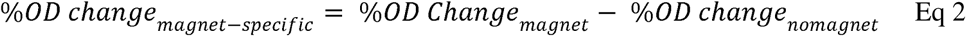

The resulting magnet-specific OD_600_ change was used as a semiquantitative measure of magnetically responsive cell-associated material and compared with total iron and oxalate-soluble iron measurements. All the OD_600_ measurements were performed with Molecular devices quick drop spectrophotometer.

### Western blotting

The spatial localization of MagA protein with polyHis tags in the MagA cells was identified by western blotting. To increase cell mass, both wild type and MagA cells were initially grown in the modified AFM3 medium with 0.1% (w/v) dispersed sulfur. The harvested cells were lysed using a FastBreak Cell Lysis Reagent for 30 min at room temperature under shaking. While both cell lysates and pellets were collected, the pellets were reacted with methanol (1:1, v/v) then heated at 50 LJ for 10 min to recover insoluble material-associated protein^27^. Additionally, the magnetosomes were purified from the MagA cells. Briefly, the magnetosomes were isolated using microfuge tube magnetic separation rack (Permagen, USA) for 2 h, then repeated discarding supernatant and resuspension the pellets in 10 mM Tris-HCl (pH 8.0) for 10 times. The final pellet was resuspended in 10 mM Tris-HCl with 1% SDS for 3 h^28^. The solution was spun down at 17,000 g for 30 min and the pellet was resuspended in 10 mM Tris-HCl. This suspension was loaded on a disposable column containing 1 mL of HisPurTM Ni-NTA resin which was equilibrated with loading buffer (20 mM Trizma, 300 mM NaCl; pH 7.4) before sample loading. The polyHis-tagged MagA protein was eluted in 2 mL of buffer containing (20 mM Trizma, 300 mM NaCl, 250 mM imidazole; pH 7.4). The eluent was collected concentrated in an Amicon® Ultra Centrifugal Filter, 30 kDa filter unit.

Both crude cell lysates and solid recovered fraction (untreated pellets extracted in methanol and purified magnetosomes) were loaded into Novex NuPAGE 4-12% Bis-Tris gels (Thermo Fisher Scientific, Waltham, MA), and the proteins were transferred onto a Immobilon®-FL PVDF membrane, using a traditional wet-transfer system. Following transfer, the membrane was blocked with 3% nonfat dry milk in TBST for 1 h at room temperature with gentle agitation. The membrane was then incubated with mouse anti-6x-His tag antibody (MA1-135; Thermo Fisher Scientific) diluted 1:1000 in blocking buffer overnight at 4 °C. After primary antibody incubation, the membrane was washed three times with TBST for 10 min each and incubated with goat anti-mouse IgG-HRP conjugate (Immun-Star GAM-HRP, Bio-Rad, #1705047) diluted 1:5000 in blocking buffer for 1 h at room temperature. The membrane was washed again three times with TBST for 10 min each, and chemiluminescent detection was performed using SuperSignal West Dura Extended Duration Substrate (Thermo Fisher Scientific, #34076). The blot was visualized using a Gel Doc XR+ Imager (Bio-Rad, Hercules, CA).

### Bioleaching experiments

Bioleaching experiments of pyrite and chalcopyrite were conducted in 14 mL Falcon Round-Bottom tubes with working volumes of 10 mL. The basal salt solution (without iron) of the modified AFM3 growth medium was used as the bioleaching medium. The wild type and engineered cells with overexpression and knockdown MagA protein were inoculated at the initial cell density of 8.3×10^7^ cells/mL (OD_600_ = 0.01). The bioleaching was performed in the absence and presence of an applied magnetic field, in which the same neodymium magnet was attached at the side of Falcon tube (Fig. S2). The pulp densities of pyrite or chalcopyrite of 1.0% (w/v) were used. All experimental conditions were performed in triplicate and incubated at 30 and 140 rpm. Supernatant samples (200 µL) were removed and filtered through a 0.2 µm pore size PVDF filter (Thermo Fisher Scientific, Waltham, MA) and subjected to AAS to measure dissolved Fe and Cu concentrations. Periodically, 200 µL aliquots were removed and diluted to measure the concentration of dissolved iron or copper by AAS. The effects of water evaporation and sample removal on the culture volume were compensated by adding distilled water to the 10 mL marking, and pH was not adjusted as it was maintained below 2 for all tested conditions throughout the experiments After bioleaching experiments, biofilm formation and cell attachment to minerals were analyzed by crystal violet assay and fluorescence in-situ hybridization (FISH), as previously described^20^.

### Other analytical methods

The OD_600_ of the cells was measured using a GENESYS 10S UV-Vis spectrophotometer. Iron and copper concentrations were measured by an iCE3300 atomic absorption spectrophotometry (Thermo Fisher Scientific, MA). Biofilm formation was measured following the previous description^20, 29^. FISH was carried out with a fluorescent oligonucleotide probe labeled with CY3 specific for *A. ferrooxidans* and ^29^ observed under Olympus Fluoview FV1000 confocal laser scanning microscope.

Statistical significance of data was evaluated by p value below 0.05 through analysis of variance (ANOVA) using Excel or PRISM.

## Results

### Enhanced magnetosome formation

The *A. ferrooxidans* genes AFE_1968 and AFE_0465 are homologous to the magnetosome-associated *magA* and *mamB* proteins^10^ and are predicted to be involved in iron transport. These genes were targeted for plasmid-driven overexpression via the tac promoter^4^ and CRISPRi-mediated knockdown using the dCas12a protein^17^. The engineered *A. ferrooxidans* cells with constitutive overexpression of the genes showed a 42-47% increase in transcriptional expression of the targeted genes, while the cells with the CRISPRi knockdown showed 57-69% reduction (Fig. 1). The engineered cells with overexpression or knockdown of putative *magA* and *magB* genes were referred to as MagA, MamB, dMagA, and dMamB cells, respectively (Table S1), and the effects of these genetic modifications on magnetosome formation in *A. ferrooxidans* were investigated.

**Figure 1.**
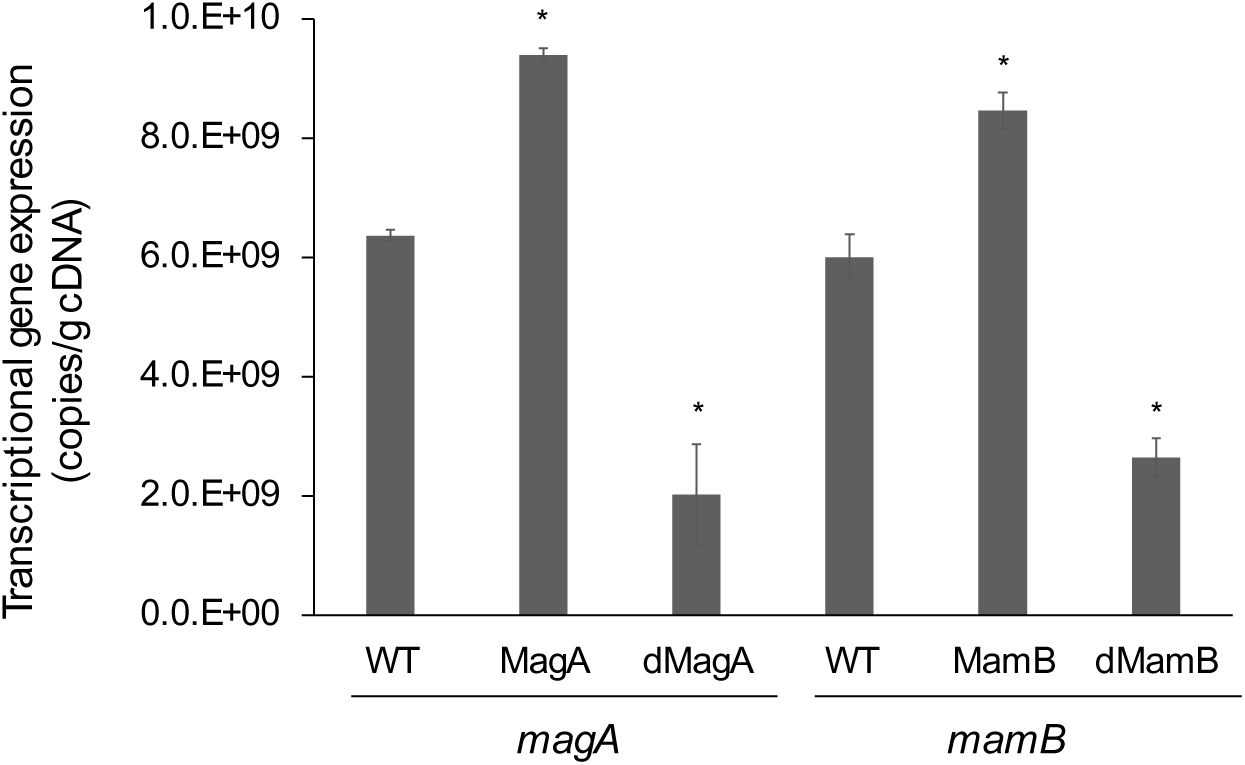
Transcriptional expression of *magA* or *mamB* genes in the wild type (WT) and engineered *A. ferrooxidans cells* with overexpression (MagA or MamB) or knockdown (dMagA or dMamB) of putative MagA and MamB proteins, respectively. Error bars are standard deviations of triplicate analyses and symbols indicate the statistical significance (*p* < 0.05) compared to the wild type cells.

After cultivation in medium optimized for magnetosome formation, total intracellular iron and oxalate-soluble iron were measured in wild type and engineered cells. The oxalate-soluble iron fraction followed a similar trend to total intracellular iron across the tested strains, suggesting enrichment of an oxalate-extractable iron phase. However, because oxalate extraction may also solubilize other iron oxide species, this measurement was interpreted as an operational proxy for magnetite-like iron rather than as a definitive measure of magnetosome-bound magnetite. When the *magA* gene expression was modulated, the MagA and dMagA cells showed 5% and 0.3% of oxalate-extractable iron based on dry cell weight, which were 2.4-fold higher and 6-fold lower than the wild type cells (Fig. 2A). This correlated with the number of magnetosomes observed under transmission electron microscopy (TEM). As compared to the wild type (1–3 smaller particles per cell) and dMagA cells (0–2 smaller particles per cell), the MagA cells had more numerous (2-4) and larger magnetosome particles (Fig. 2B, Fig S3). On the other hand, the MamB and dMamB cells showed comparable total iron and magnetite contents with the wild type cells (Fig. 2A). These results indicate that the *magA* gene (AFE_1968), encodes for a MagA protein, has a regulatory role in magnetosome formation in *A. ferrooxidans*, as opposed to the putative *mamB* gene (AFE_0465) which seemed to have little effect. Therefore, subsequent experiments focused on the expression of the *magA* gene, in comparison with the wild type cells.

**Figure 2.**
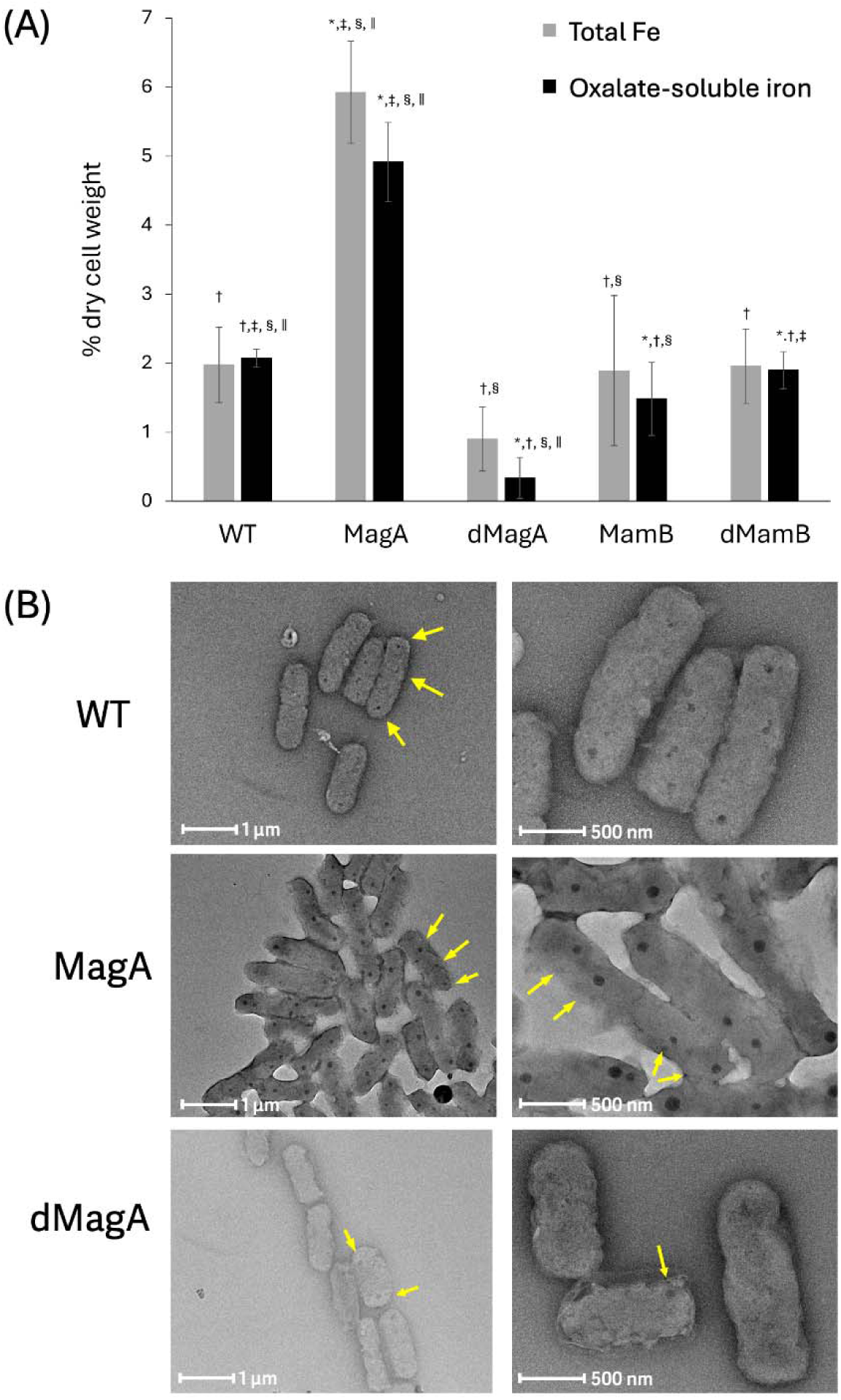
(A) Intracellular total iron and oxalate-soluble Fe concentrations in the wild type (WT) and engineered *A. ferrooxidans* cells with overexpression (MagA or MamB) and knockdown (dMagA or dMamB) of putative *magA* and *mamB* genes. (B) TEM characterization of magnetosomes in the wild type and engineered cells. All cells were incubated for 60 h prior to harvest for the analysis. In panel A, error bars indicate the standard deviations of triplicate analyses, and symbols indicate the statistical significances (*p* < 0.05; *, compared to WT; †, compared to MagA; ‡ compared to dMagA; §, compared to MamB;, compared to dMamB). In panel B, magnetosomes are indicated by yellow arrows. Additional TEM images can be found in Fig S3.

The recombinant proteins were expressed with C-terminal polyhistidine tags, which allow for Western blot detection. The *magA* gene is thought encode a proton-driven Fe (II) antiporter protein and it likely serves an important role in iron uptake^13, 30^, the tagged MagA protein was detected in the insoluble fractions of the MagA cells. The magnetosomes were further purified from the MagA cell line and the tagged MagA protein was found to be associated with the magnetosomes in the cells (Fig. 3A). This association is consistent with a possible role for MagA in magnetosome-associated iron handling or magnetosome-like particle formation. On the other hand, it was also observed that the MagA cells both oxidized Fe^2+^ and grew more rapidly than the wild type cells, which was not observed when the cells were grown on sulfur (Fig. 3B), suggesting that the effect of MagA expression is linked to iron-dependent growth conditions.

**Figure 3.**
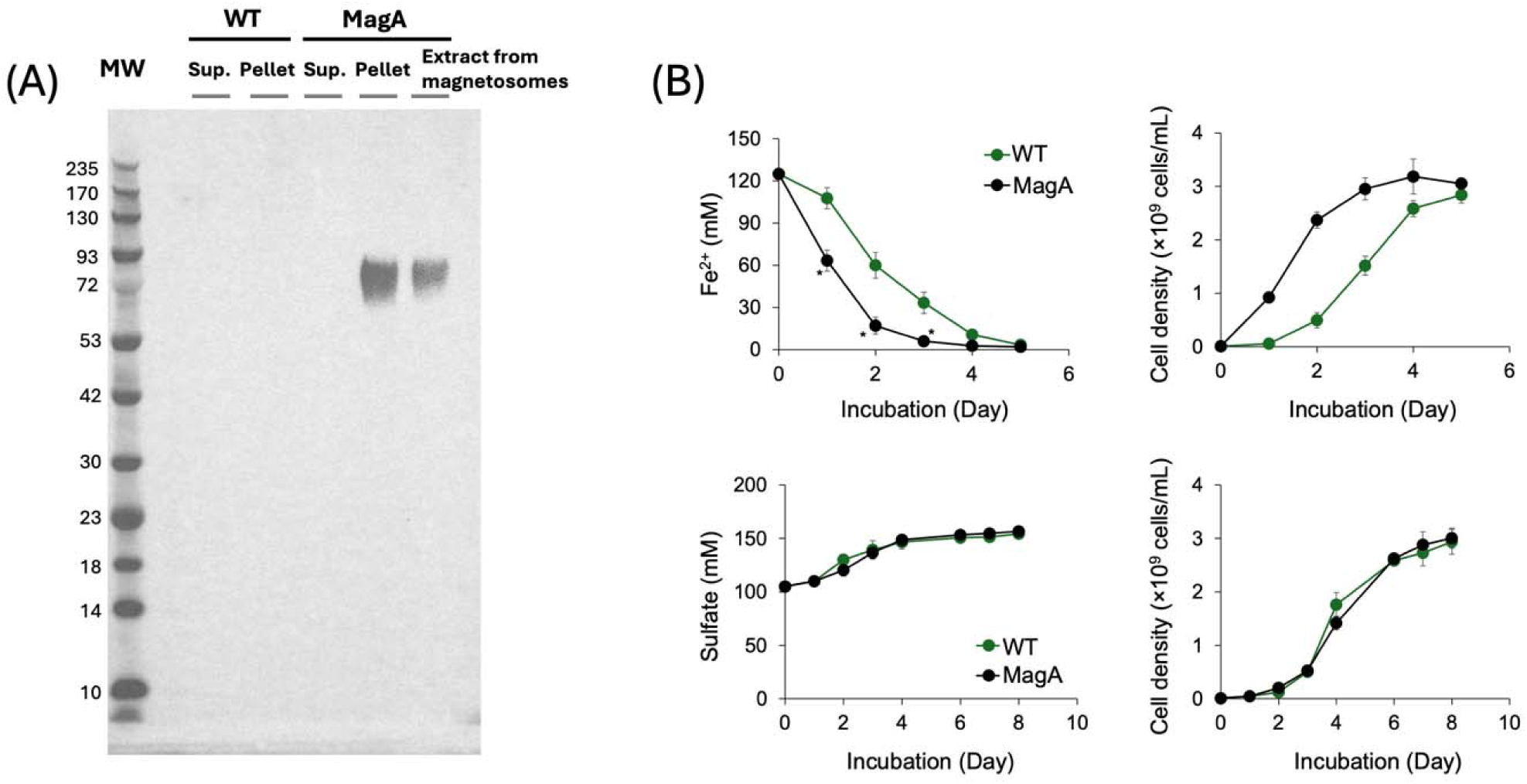
(A) Western blot assays of cell lysates from WT and MagA cells, pellets of MagA cells, and purified magnetosomes (Mag) from the MagA cells, using polyhistidine tags. His-tagged MagA has a predicted molecular weight of ∼71.3 kDa, and a corresponding band near this size was detected only in the pellet and magnetosome-enriched extract fractions from MagA cells. (B) Substrate utilizations and growth curves of the wild type (WT) and engineered *A. ferrooxidans* cells with overexpression of putative *magA* gene (MagA), under iron or sulfur as sole substrates. Error bars indicate the standard deviations of triplicate analyses and symbols show statistical significances (*p* <0.05) compared to the wild type.

Given the magnetic characteristic of magnetosomes, it seems likely that *A. ferrooxidans* would respond to, and potentially be capable of orienting towards, magnetic fields. However, the weak native magnetotaxis of *A. ferrooxidans* compared MTB makes it difficult to observe the magnetophoresis effect^10, 31^. We explored whether the increased magnetosome formation in the MagA cells could amplify this response by exposing cells to magnetic forces.

To this end, a modified whole-cell magnetic-response assay was used to evaluate magnetically responsive cell-associated material in WT, MagA, MamB, and dMagA samples (Fig. 4). Cell suspensions were normalized to OD_600_ = 0.4 and incubated for 10 min in either a cuvette fitted with two external neodymium magnets or a matched no-magnet control cuvette (Fig. 4A, Fig. S1). In no-magnet controls, OD_600_ changed by no more than 0.5% over the 10 min assay period, indicating minimal passive settling under these conditions.

**Figure 4.**
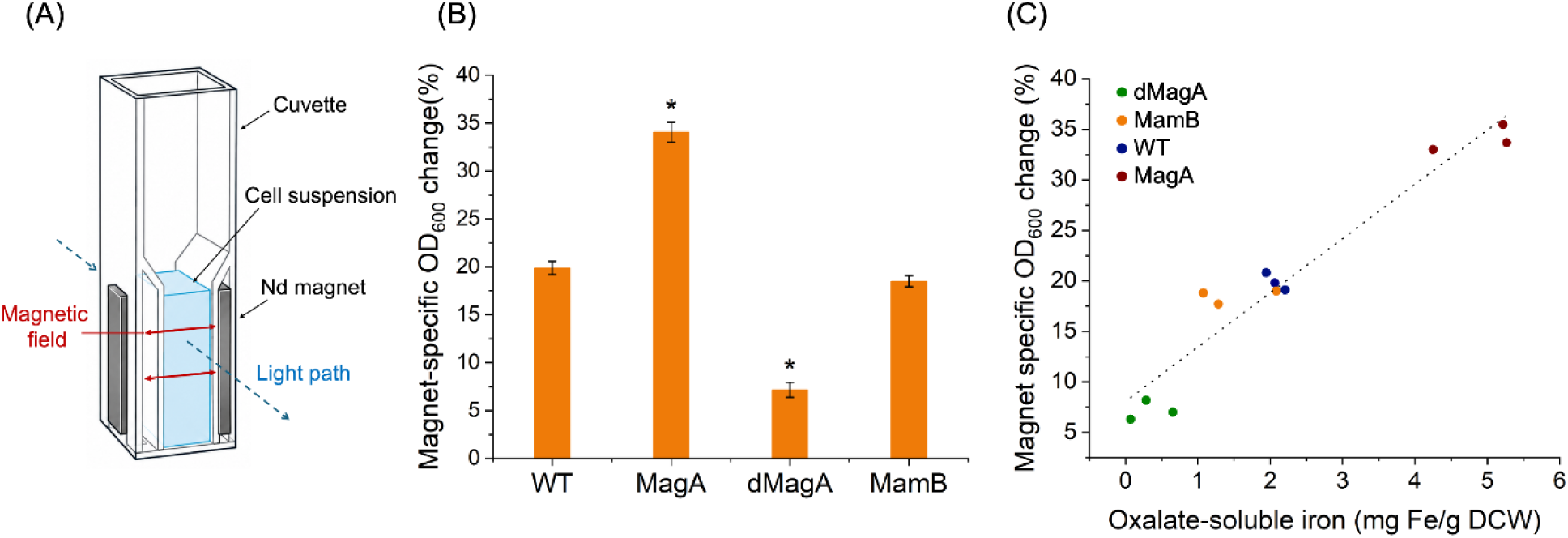
Modified whole-cell magnetic-response assay to assess magnetically responsive cell-associated iron. (A) Schematic of the modified cuvette-based magnetic-response assay. Cell suspensions were placed in a cuvette fitted with two external neodymium magnets positioned on opposite side walls (Fig S1). (B) Magnet-specific OD_600_ change measured after 10 min magnetic exposure for WT, MagA, dMagA, and MamB cells. Bars represent mean ± SD from three replicates, and symbols indicate statistical significance (p < 0.05) compared to wild type cells. (C) Relationship between oxalate-soluble iron content and magnet-specific OD_600_ change across the tested strains. Individual replicate values are shown, with the dotted line indicating the trend between oxalate-soluble iron and magnetic response.

Magnet exposure resulted in strain-dependent OD_600_ depletion (Fig. 4B). The dMagA sample showed the lowest magnet-specific OD_600_ change (7.3 ± 0.9%), while wild type and MamB samples showed intermediate responses (20 ± 1% and 19 ± 1%, respectively). The MagA sample showed the highest magnet-specific OD_600_ change (35 ± 1%). Thus, the magnitude of magnet-induced OD_600_ depletion increased from dMagA to wild type/MamB and was highest in the MagA sample.

When magnet-specific OD_600_ change was plotted against oxalate-soluble iron content (Fig. 4C), the samples showed a positive association between magnetic response and the oxalate-soluble iron fraction. This relationship was observed across the replicates analyzed for dMagA, wild type, MamB, and MagA.

### Effects of enhanced magnetosomes on bioleaching

The role of magnetosomes in metal bioleaching has not been fully elucidated. The MagA and dMagA cells provide a range of magnetosome concentrations and we examined the effect of magnetosome loading on the bioleaching of weakly paramagnetic pyrite (FeS_2_) ^32^ in the absence or presence of magnetic fields (Fig. 5), and this was compared with non-magnetic iron sulfide, chalcopyrite (CuFeS_2_; antiferromagnetic^33^). At the end of the pyrite bioleaching, the final iron concentrations measured in the wild type and dMagA cell cultures across both magnetic conditions were 200–250 and 190–210 mg/L, respectively, showing no significant effect of the applied magnetic field on the soluble iron concentration (Fig. 5A) or the fold change in pyrite dissolution relative to the corresponding wild type controls (Fig. S4A). By contrast, the MagA cells showed an increased dissolved iron concentration of ∼1,000 mg/L in the absence of magnetic fields, and this was further increased to ∼1,200 mg/L under application of an additional external magnetic field. These increased iron concentrations corresponded to 1.5– and 1.8–fold increases in pyrite dissolution, which were significantly higher than the corresponding wild type controls in the absence and presence of applied magnetic field, respectively (Fig. S4A). In contrast, the final concentration of copper dissolved from the chalcopyrite sample tested was comparable among the engineered cell lines, which were comparatively lower than the corresponding wild type controls regardless of the presence of an applied magnetic field (Fig. 5B), though the chalcopyrite dissolution was only significantly reduced relative to the wild type control for both MagA and dMagA in the absence of the applied magnetic field (Fig. S4B). These experiments were designed to compare bioleaching performance among cell-containing conditions under identical acidic medium, mineral loading, incubation time, and magnetic-field conditions. Therefore, the data are interpreted as relative differences among wild type, MagA, and dMagA conditions rather than as an absolute quantification of biological leaching above abiotic acidic dissolution.

**Figure 5.**
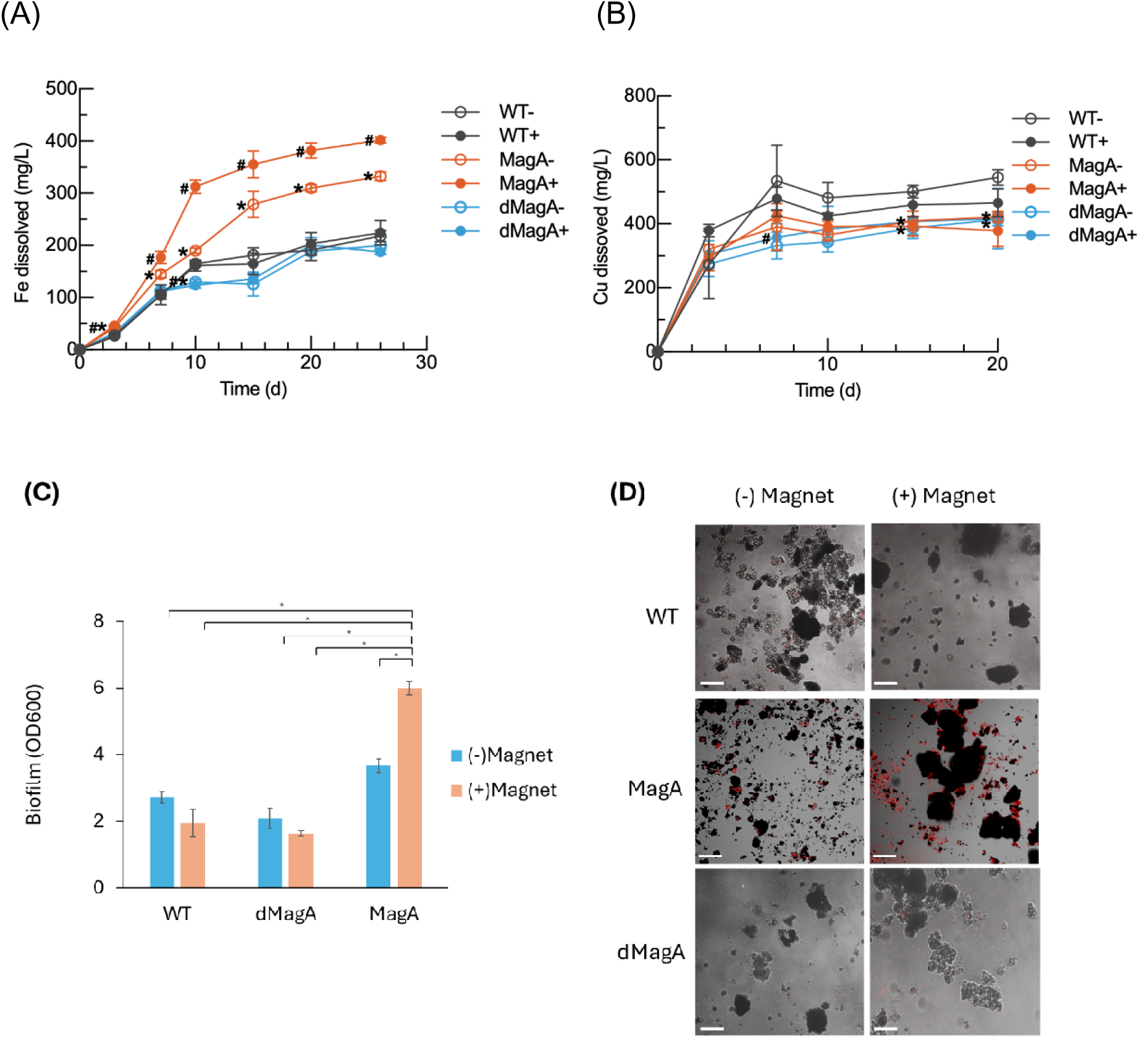
Time-course measurements of (A) iron and (B) copper dissolution from pyrite and chalcopyrite, respectively, under the conditions with the wild type (WT) and engineered *A. ferrooxidans* cells with overexpression (MagA) and knockdown (dMagA) of putative MagA protein, either in the absence (–) or presence (+) of magnetic fields. Symbols indicate the statistical significance of the adjusted comparison to the corresponding wild type condition using two-way ANOVA with Dunnett’s multiple comparisons test (*P < 0.05 relative to WT–; #P < 0.05 relative to WT+). (C) Biofilm formation (* indicates the statistical significance (*p* < 0.05) between the different conditions) and (D) FISH measurement after the bioleaching of pyrite. In panel D, *A. ferrooxidans* cells are stained with CY3 and the bars represent 5 mm.

In addition to enhanced bioleaching, higher levels of biofilm formation by the MagA cells were observed during the bioleaching of pyrite, particularly under magnetic fields, as compared to the wild type and dMagA cells (Fig. 5C). The enhanced adherence to the pyrite surface by the MagA cells suggests the magnetosomes enable the sensing of the paramagnetism of the pyrite which is enhanced in the presence of the external magnetic field (Fig. 5D).

## Discussion

*A. ferrooxidans* is an important microbe for industrial metal bioleaching and there is interest in developing it as a platform for other applications^4^ including biocorrosion^34, 35^, biochemical production^36, 37^, and the production of biominerals^13, 30^. *A. ferrooxidans* are known to create magnetosomes, and it has been suggested that these cells could be superior for magnetosome manufacturing as compared to other MTB. We set out to explore the role of the *magA* and *mamB* homologs in *A. ferrooxidans* by both constitutively overexpressing the genes and knocking down the expression of the genes using dCas12a mediated CRISPRi^17^.

Of the two genes investigated, we found that only *magA* gene appears to affect magnetosome synthesis in *A. ferrooxidans* (Fig. 2). In MTB, the MagA protein has been predicted to be a proton-driven Fe (II) antiporter protein and it has been proposed to serve an important role in iron uptake, which is required for magnetosome synthesis^13, 30^. The MagA from *Magnetospirillum magneticum* AMB-1 has been shown to enable iron uptake in extracellular vesicles ^38^ and in mammalian cells^39^. On the other hand, *magA* deletions in both *Magnetospirillum magneticum* AMB-1 and *Magnetospirillum gryphiswaldense* MSR-1 had no impact on growth or magnetosome formation in either strain^40^. Here we show that the overexpression of MagA protein not only increases the number of magnetosomes (Fig 2), but also led to an increased response of the cells to a magnetic field (Fig. 4).

A comparison of the *A. ferrooxidans* AFE_1968 MagA-like homolog with representative MagA proteins from magnetotactic bacteria is provided in Table S4, highlighting that MagA-related proteins have been associated with magnetosome biology but may have species-dependent functional importance. The MagA protein was found to localize with the magnetosomes (Fig 3A) suggesting it may be involved in the formation of the magnetosomes. Interestingly, its role in iron uptake was also confirmed via the observation that the MagA overexpressing cells oxidized iron and grew faster than the wild type cells (Fig 3B). These results confirm that genetic engineering can be used to increase magnetosome formation in *A. ferrooxidans*, and these results may be further enhanced as these cells are developed as a platform for nanomaterial biosynthesis.

We initially sought to perform a C_mag_-type light-scattering assay similar to the method described by Schüler, Uhl, and Bäuerlein, in which magnetic cells are exposed to magnetic fields parallel and perpendicular to the light path and the resulting change in light scattering is used as a proxy for cellular magnetism^26^. In that assay, magnetosome-containing *Magnetospirillum gryphiswaldense* cells align in an applied magnetic field because organized intracellular magnetite chains generate a cellular magnetic dipole. However, our spectrophotometer geometry did not permit controlled application of a magnetic field parallel to the light beam. In addition, preliminary trials did not show a reliable OD_600_ change under the original alignment-style configuration, which may reflect the weaker and less canonical magnetic behavior expected for *A. ferrooxidans* compared with magnetotactic bacteria such as *M. gryphiswaldense*.

Therefore, we developed a modified single-orientation whole-cell magnetic-response assay and this was used as an orthogonal, semiquantitative readout of magnetically responsive cell-associated material. The rationale is that oxalate-dissolved AAS measurements may detect iron released from magnetite but may also include other ferric oxides or nonspecific cellular iron deposits. Such nonspecific iron pools may contribute to total iron content without producing a magnetic response. By contrast, magnetically responsive cell-associated particles should cause magnet-dependent OD_600_ depletion under a fixed field geometry. The observed strain-dependent magnetic response, together with its positive association with oxalate-soluble iron, supports the interpretation that at least part of the oxalate-soluble fraction reflects magnetically responsive iron-containing material. This supports the strain-dependent magnetic response however it has important limitations. It is not a formal C_mag_ measurement because only a single magnetic-field orientation was tested, and it does not directly quantify magnetosome number. It also does not chemically prove that the responsive material is magnetite, nor does it demonstrate that the particles are membrane-bound intracellular magnetosomes. Rather, the assay provides functional evidence for magnetically responsive cell-associated iron particles, which complements total iron and oxalate-soluble iron measurements. A definitive assignment of magnetite or magnetosome structure would require additional mineralogical or ultrastructural analyses.

The enhanced magnetic sensitivity exhibited by the MagA cells affected the bioleaching of pyrite, which is weakly paramagnetic (Fig. 5). Cell attachment is an essential step for bioleaching of minerals ^20^, and the enhanced biofilm formation and cell adherence to pyrite correlated with increased iron solubilization (Fig. 5A, Fig S4A). However, these measurements do not distinguish whether the magnetic field directly enhances cell-mineral attachment or promotes local enrichment/aggregation of magnetically responsive cells, which could secondarily increase mineral contact and biofilm formation. Therefore, we interpret the increased biofilm signal as enhanced mineral-associated cell accumulation under magnetic-field conditions. These results were not observed in bioleaching of non-magnetic chalcopyrite (Fig. 5B, Fig S4B), and thus it suggests positive interactions among the engineered cells, magnetic minerals, and magnetic fields found in the environment. The improved pyrite bioleaching results also suggest the potential application of the MagA cells for resource recovery from various magnetic sources, including iron minerals and magnetized electronic waste^41-43^.

Although these results support a functional role for MagA in magnetosome-associated iron accumulation and magnetic phenotype in *A. ferrooxidans*, the direct molecular mechanism remains unresolved. The present data do not establish whether MagA directly transports Fe(II), whether transport is proton-coupled, or whether MagA functions independently or through interactions with MamB homologs or other magnetosome-associated membrane proteins. Direct mechanistic studies will require biochemical reconstitution of purified MagA into defined membrane systems. Future work using MagA-containing proteoliposomes^44-46^, Fe(II) uptake kinetics, pH-gradient or membrane-potential-dependent transport assays, and protein-interaction approaches such as co-immunoprecipitation or crosslinking-mass spectrometry will be important for defining the molecular basis of MagA-dependent iron handling during magnetosome formation.

## Conclusion

The overexpression and the CRISPRi knockdown of the *magA* gene in *A. ferrooxidans* were used to modulate the formation of magnetosomes within these cells. *A. ferrooxidans* is being developed as chassis strain for novel biotechnology applications, and these results demonstrate the potential for engineering for nanomaterial biosynthesis. Unexpectedly, the increased magnetosomes led to enhanced magnetic sensitivity by the cells leading to augmented pyrite bioleaching which could be further enhanced by a magnetic field. These results confirm an important role for these structures in *A. ferrooxidans* cellular physiology and demonstrate that sulfide mineral bioleaching can be manipulated via magnetosome engineering.

## Acknowledgements

This work was supported by ARPA-E grant DE-AR0001340 from the US Department of Energy. The authors would like to thank Dr. Hannah Zurier and Mallori Herishko for technical assistance.

## Author Contributions

H.J. and S.B. conceived and designed the research. H.J. and S.A. performed the experiments. S.A. and Z.S. performed the TEM experiments. H.J., S.A. and S.B. wrote and edited the manuscript.

## Supporting Information

Materials in supporting information include 4 Tables and 4 Figures. Supporting Tables include Plasmids and strains, Primer sequences, Sequences of guide RNAs, Comparison of MagA-like proteins. Supporting Figures include Images of cuvette setup for the whole-cell magnetic-response assay, Images of bioleaching conditions in the presence of a magnet, Additional TEM images of magnetosomes, Fold differences in iron and copper dissolution.

