## Supporting Information for "Overexpression of *magA* in *Acidithiobacillus ferrooxidans* increases magnetosome production and pyrite bioleaching"

Supporting information includes:

4 Tables: Plasmids and strains, Primer sequences, Sequences of guide RNAs, Comparison of MagA-like proteins

4 Figures: Images of cuvette setup for the whole-cell magnetic-response assay, Images of bioleaching conditions in the presence of a magnet, Additional TEM images of magnetosomes, Fold differences in iron and copper dissolution.

**Table S1.** Plasmids and strains used in this study.

| Strains or Plasmids | Description | Source or Reference |
|---|---|---|
| ***Strains*** | | |
| ***E. coli*** **DH5α** | *fhuA2Δ(argF-lacZ)U169 phoA glnV44 Φ80Δ(lacZ)M15 gyrA96 recA1 relA1 endA1 thi-1 hsdR17* | NEB |
| ***E. coli*** **DH10β** | *Δ(ara-leu) 7697 araD139 fhuA ΔlacX74 galK16 galE15 e14- ϕ80dlacZΔM15 recA1 relA1 endA1 nupG rpsL (*$Str^R$*) rph spoT1 Δ(mrr-hsdRMS-mcrBC)* | NEB |
| ***E. coli*** **S17-1 (ATCC 47055)** | *recA pro hsdR* RP4-2-Tc::Mu-Km::Tn7 integrated into the chromosome | ATCC |
| ***A. ferrooxidans*** **(ATCC 23270)** | Type strain | ATCC |
| **MagA** | ATCC 23270 with pMagA | This study |
| **MamB** | ATCC 23270 with pMamB | This study |
| **dMagA** | ATCC 23270 with pdMagA | This study |
| **dMamB** | ATCC 23270 with pdMamB | This study |
| ***Plasmids*** | | |
| **pYI39** | pJRD215 empty vector with *tac* promoter and rrnB terminator | [1] |
| **pJRD_dCas12a** | dCas12a/CRISPRi constructed on pJRD215 vector | [2] |

| | | |
|---|---|---|
| **pMagA** | pYI39 with putative MagA (AFE_1968) with polyHis tags | This study |
| **pMamB** | pYI39 with putative MamB (AFE_0465) with polyHis tags | This study |
| **pdMagA** | pJRD_dCas12a for *magA* knockdown | This study |
| **pdMamB** | pJRD_dCas12a for *mamB* knockdown | This study |

The nucleotide sequences of pMagA, pMamB, pdMagA, and pdMamB are deposited in Genbank with accession numbers of PP597399–597402.

**Table S2**. Primer sequences used in this study.

| | | Sequence (5'→3') |
|---|---|---|
| *Cloning* | | |
| pMagA | F | ATCCCCTTAGAATTCGTTCTGGTACATGCAAGATCACGGCCTGTT |
| | R | TTGTCAGTGGTGATGGTGATGATGGTTAACTTCCTCCTCCACCCGCTTCCTGTACTTTAT |
| pMamB | F | ATCCCCTTAGAATTCGTTCTGGTACATGGCGGAAAGCATCAGCCG |
| | R | TTGTCAGTGGTGATGGTGATGATGGTTATGGGGCGCCGGATGACA |
| *Site-directed mutagenesis* | | |
| pdMagA | F | GTTGTAGATCGATCAGTACGGTGCTGCGTCAAATAAAACGAAAGGCTC |
| | R | AGTAGAAATTTAATGACTACAGCCCGTGCAATCTACAACAGTAGAAATTATTTAAAG |
| pdMamB | F | GTTGTAGATTGGACGCTCTATATTGCCGCCAAATAAAACGAAAGGCTC |
| | R | AGTAGAAATTTCGCTAGCCTGCAGGAGACGATCTACAACAGTAGAAATTATTTAAAG |
| *dPCR* | | |
| *magA* | F | TCGGACTTGCGTGGATTAC |
| | R | GATATCCGCTTCCACCTGATAG |
| *mamB* | F | GCCACCAAAGAAACCCTCTAT |
| | R | AGACGATGACGCTGGAAATG |

**Table S3**. Sequences of guide RNAs (gRNA) used in this study.

| Target gene | Strand | Sequence (5'→3') | PAM |
|---|---|---|---|
| *magA* | + | TGCACGGGCTGTAGTCATTA | TTTC |
| | − | CGATCAGTACGGTGCTGCGT | TTTC |
| *mamB* | − | CGTCTCCTGCAGGCTAGCGA | TTTC |
| | − | TGGACGCTCTATATTGCCGC | TTTC |

**Table S4:** Comparison of the *A. ferrooxidans* MagA-like homolog with representative MagA proteins from magnetotactic bacteria.

| Organism | Protein / locus | Annotation | Reported role | Reported phenotype | Reference |
|---|---|---|---|---|---|
| *A. ferrooxidans* ATCC 23270 | AFE_1968 | Potassium-efflux-system / cation antiporter-like protein | Candidate MagA-like transporter associated with magnetosome-like particle formation | Overexpression increased oxalate-soluble iron, TEM-visible particles, and magnetic response; knockdown reduced these phenotypes | This study; & Liu et al.[3] |
| *Magnetospirillum magneticum* AMB-1 | MagA | Iron transport-related membrane protein / MagA | Originally proposed to participate in iron transport for magnetosome formation | Early studies suggested a transport-related role | Nakamura et al.[4] |
| *Magnetospirillum gryphiswaldense* MSR-1 | MagA homolog | MagA-like membrane protein | Tested in relation to magnetosome formation | Deletion or comparative studies suggested little or no essential contribution to magnetite biomineralization | Uebe et al.[5] |

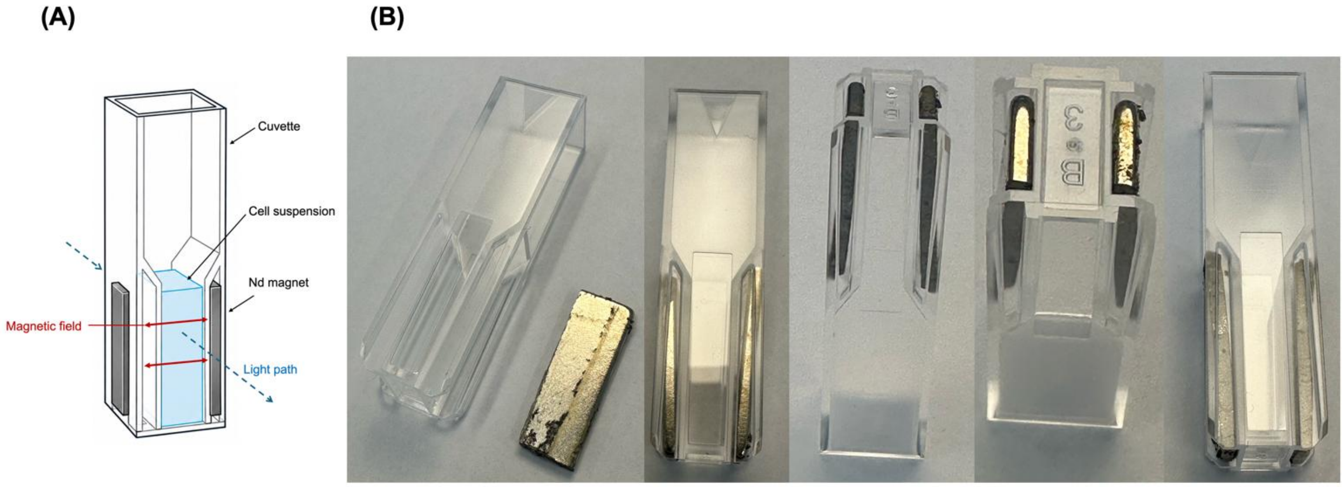


**Figure S1**. Modified cuvette setup for the whole-cell magnetic-response assay. (A) Schematic representation of the modified cuvette-based magnetic-response assay. Two neodymium magnets were positioned externally on opposite side walls of a BRAND® 1.5 mL semi-micro cuvette to generate a lateral magnetic field across the cell suspension. The magnets were placed outside the sample chamber and oriented so that the magnetic field passed across the cuvette without obstructing the spectrophotometer light path. (B) Representative photographs of the modified cuvette assembly showing the external magnet placement, and the unobstructed optical faces used for $OD_{600}$ measurements. The setup was used to measure magnet-induced $OD_{600}$ depletion as a semiquantitative readout of magnetically responsive cell-associated material.

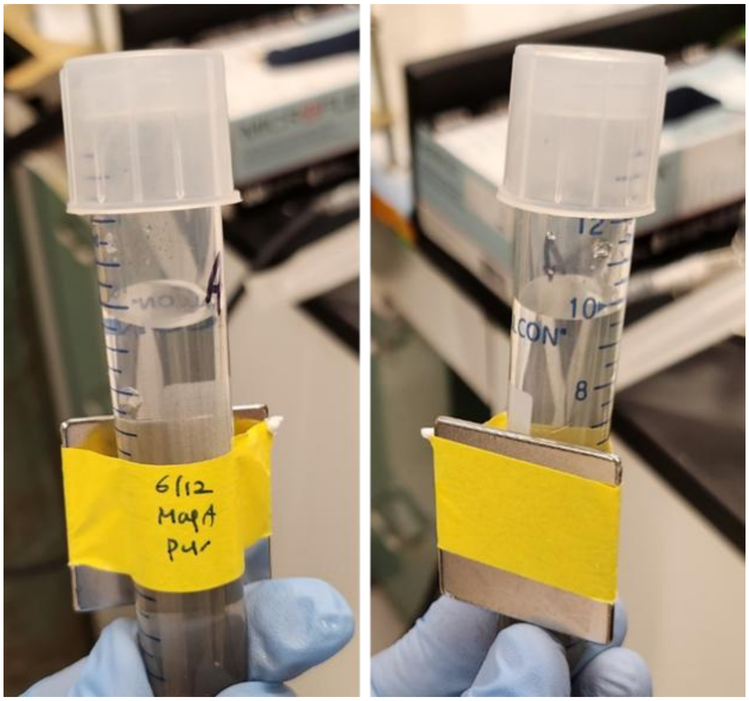

**Figure S2**. Images of bioleaching experiments in the presence of magnetite. A permanent neodymium magnet was attached to the side of 15 mL falcon tube used for the bioleaching of pyrite and chalcopyrite.

**A**

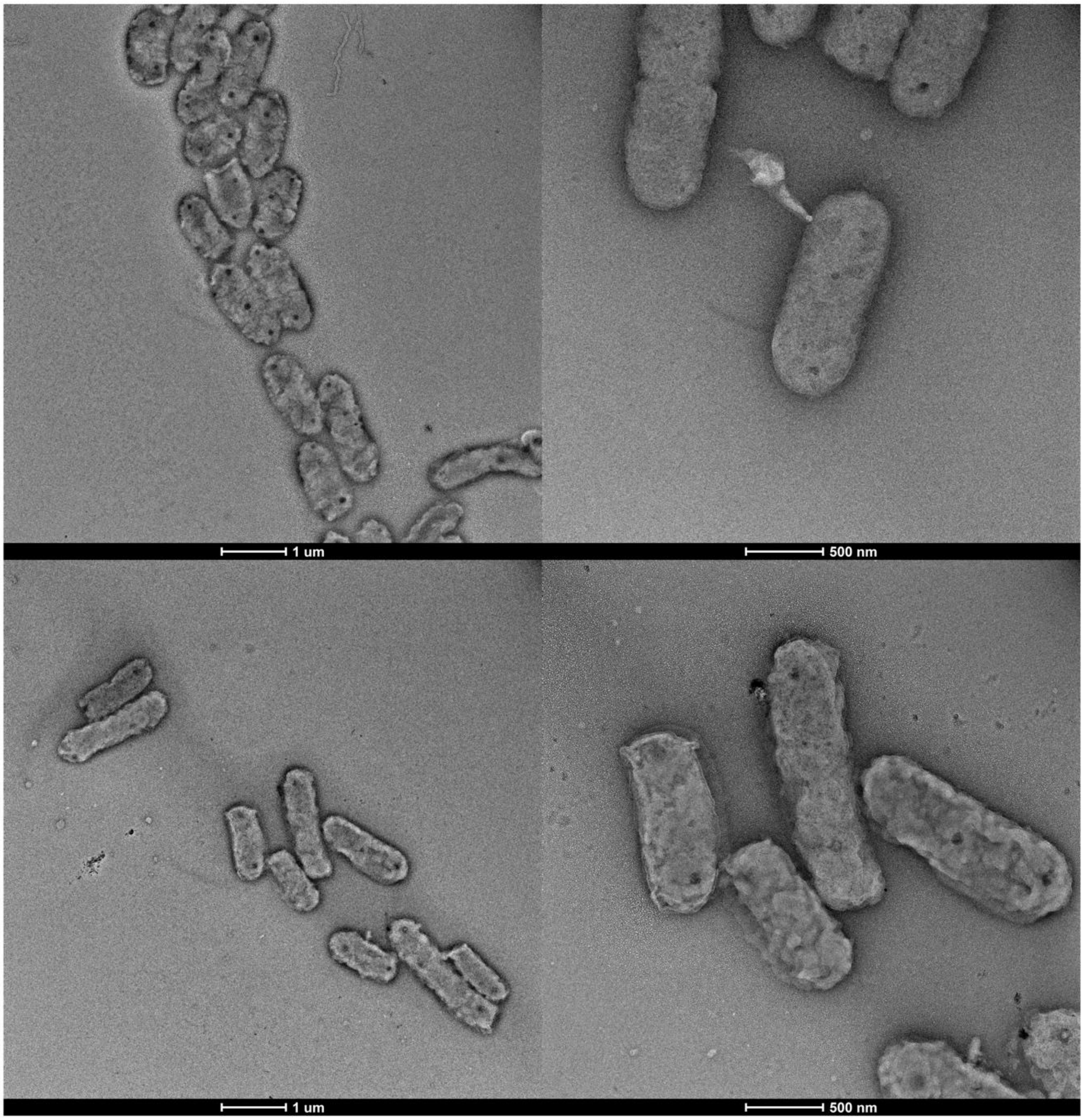

**B**

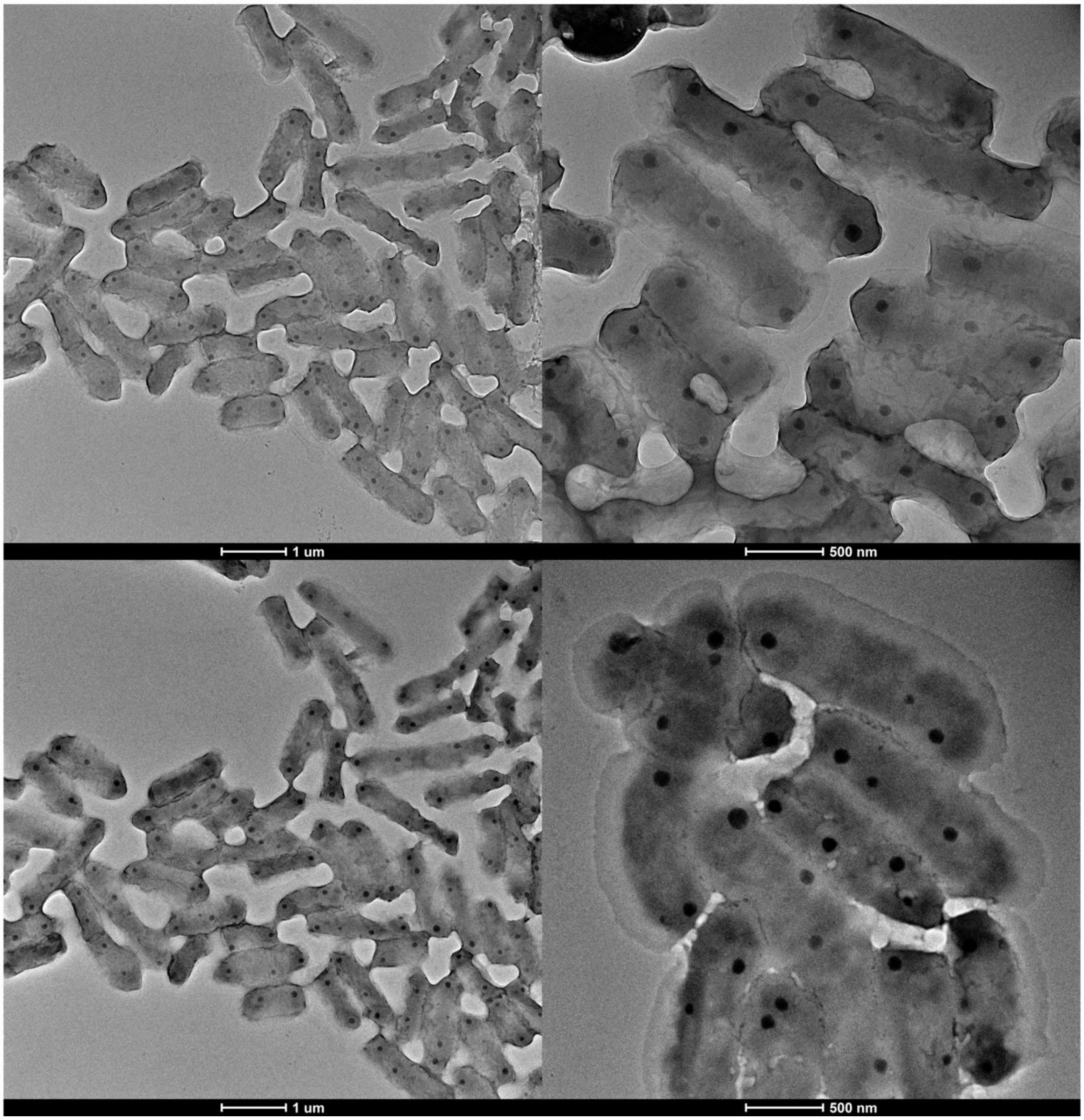

**C**

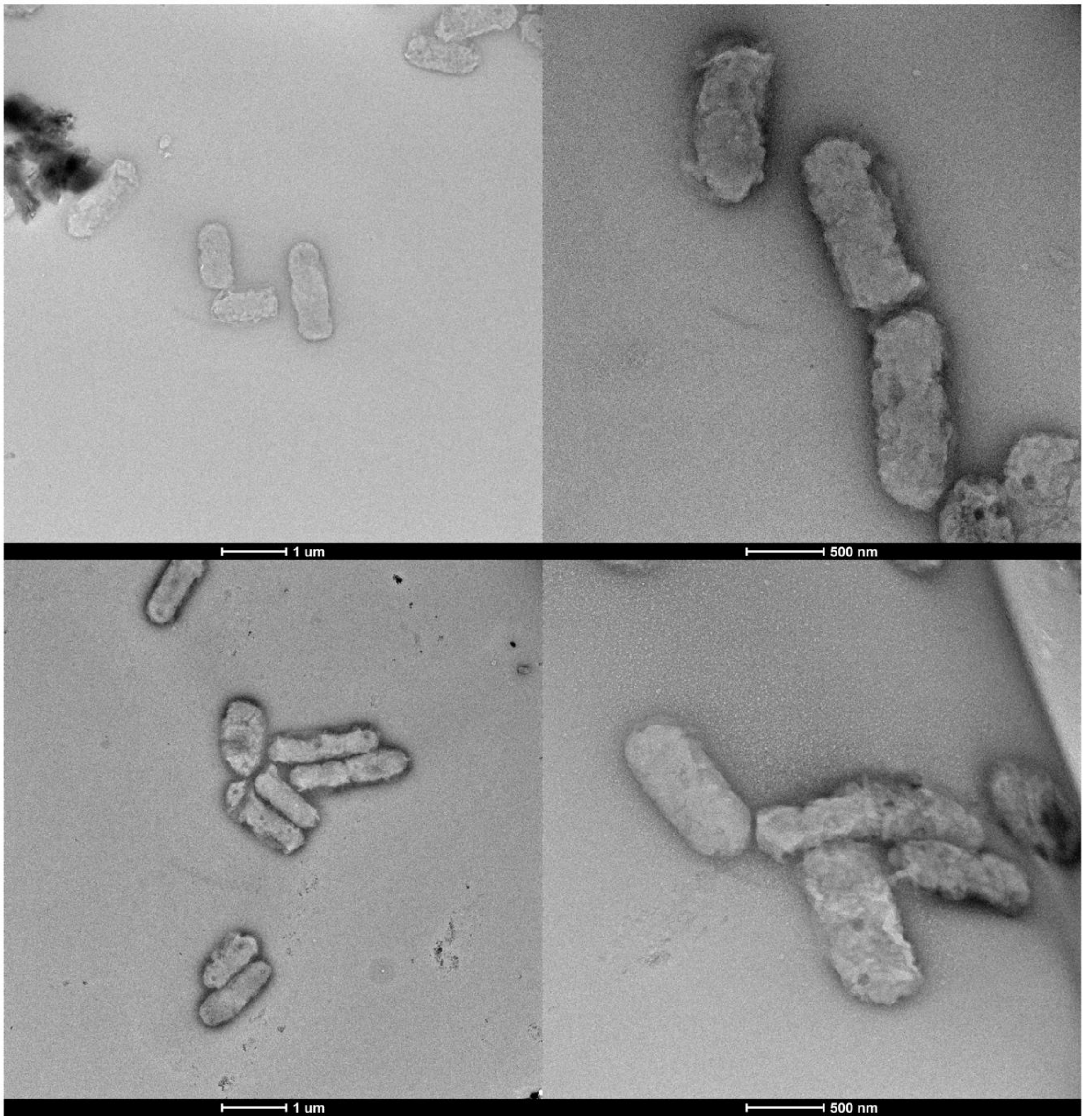


**Figure S3**. Additional TEM Images of (A) wild type cells, (B) MagA cells, and (C) dMagA cells.

A

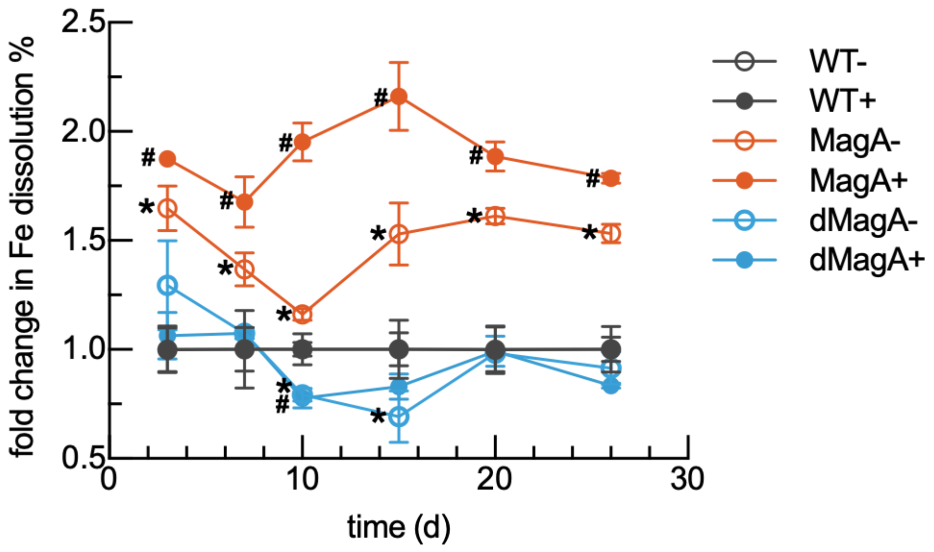


B

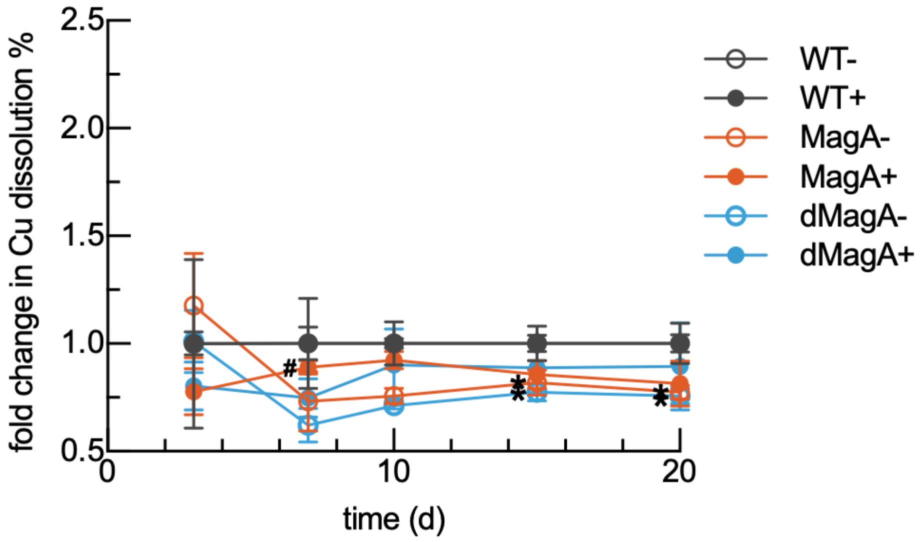


**Figure S4**. Fold changes in dissolution of iron from pyrite (A) and copper from chalcopyrite (B) using data from Figure 5. Symbols indicate the statistical significance of the adjusted comparison to the corresponding wild type condition using two-way ANOVA with Dunnett's multiple comparisons test (*P < 0.05 relative to WT–; #P < 0.05 relative to WT+).
